# Leaf chemical defences and insect herbivory in oak: accounting for canopy position unravels marked genetic relatedness effects

**DOI:** 10.1101/872549

**Authors:** Elena Valdés-Correcher, Audrey Bourdin, Santiago C. González-Martínez, Xoaquín Moreira, Andrea Galmán, Bastien Castagneyrol, Arndt Hampe

## Abstract

**Background and Aims:** Highly controlled experiments revealed that plant genetic diversity and relatedness can shape herbivore communities and patterns of herbivory. Evidence from the field is scarce and inconsistent. We assessed whether a genetic signal underlying herbivory can be detected in oak forest stands when accounting for variation at smaller (within-tree) and larger (among-stand) scales.

**Methods:** We tested relationships between tree genetic relatedness, leaf chemical defences and insect herbivory at different canopy layers in 240 trees from 15 Pedunculate oak (*Quercus robur*) forest stands and partitioned sources of variability in herbivory and defences among stands, individuals, and branches.

**Key Results:** Leaf defences, insect herbivory, and their relationship differed systematically between the upper and the lower tree canopy. When accounting for this canopy effect, the variation explained by tree genetic relatedness rose from 2.8 to 34.1 % for herbivory and from 7.1 to 13.8 % for leaf defences. The effect was driven by markedly stronger relationships in the upper canopy.

**Conclusions:** Our findings illustrate that properly accounting for other sources of variation acting at different scales can reveal potentially relevant effects of the host plant genotype on patterns of leaf chemical defences and associated insect herbivory in natural tree populations.

## Introduction

A rapidly growing number of studies have shown over the last decade that plant genetic diversity and genetic relatedness can influence herbivore communities and associated patterns of herbivory (Crutsinger *et al.* 2006; McArt and Thaler 2013; Kagiya *et al.* 2018). It has been proposed that the composition and activity of herbivore communities are heritable traits of the host plant that are partly driven by the heritability of its anti-herbivore chemical defences (Wimp *et al.* 2005; Bangert *et al.* 2006; Bustos-Segura *et al.* 2017; Jenkins and Brown 2018). Plant families vary indeed considerably in their edibility and resulting herbivore damage (Donaldson and Lindroth 2007; Fernandez-Conradi *et al.* 2017; Barker *et al.* 2018; Damestoy *et al.* 2019). However, most previous research has been performed on highly-controlled experiments (e.g. common gardens), often with juvenile plants and minimized spatial and environmental effects, settings that could lead to overemphasize the putative role of genetics in nature (Tack *et al.* 2012; Lämke and Unsicker 2018). Accordingly, more research in natural plant populations is needed for understanding to which extent genetically-based variation in plant chemical defences determines insect herbivory (Wimp *et al.* 2005; Carmona *et al.* 2011).

Diverse biological mechanisms can contribute to blur links between plant genotype, plant chemical defences and herbivory patterns in mature trees in the wild. Many secondary metabolites exhibit low heritability because their production is controlled by multiple genes and their interactions (Külheim *et al.* 2011; Büchel *et al.* 2016). Different plant parts experience different microclimates (e.g., irradiation, temperature, humidity) that can trigger extensive within-individual variation in leaf morphology and chemistry, especially along tree vertical gradients. Upper canopy leaves are typically thicker, tougher, smaller, drier and contain higher levels of chemical leaf defences than lower canopy leaves (Murakami and Wada 1997; Le Corff and Marquis 1999; Murakami *et al.* 2005; Ruhnke *et al.* 2009; De Casas *et al.* 2011; Castagneyrol *et al.* 2019). More specifically, differences in microclimate should directly affect the expression of genes that code the production of leaf chemical defences (reviewed in Lämke and Unsicker, 2018). In turn, tree vertical gradients in insect herbivory can result from differences in herbivore dispersal (e.g., flying insects concentrated in the upper canopy; Ulyshen, 2011) or herbivore exposition to predators (e.g., lower predation rates in the upper canopy; Aikens et al., 2013) that are not driven by leaf chemistry. Genotype-phenotype-herbivory associations can also be obscured at larger spatial scales owing to the non-random distribution of host plant genotypes (i.e., spatial or population genetic structure) that is widespread within and among natural plant populations as a consequence of limited effective gene flow and/or genetic drift (Hoban et al., 2016; Rellstab et al., 2015; see also Tack et al., 2012). Finally, landscape-scale patterns of herbivore abundance and diversity are well-known to be strongly influenced by resource availability and by herbivores’ spatial grain of habitat perception and use (Tack *et al.* 2010; O’Rourke and Petersen 2017; Bagchi *et al.* 2018; Valdés-Correcher *et al.* 2019). The plethora of potential confounding factors underpins that careful study designs including multiple-scale sampling are needed to thoroughly assess effects of genetically-based variation in leaf chemical defences on herbivory in natural plant populations.

This study investigated the relationships between tree genetic relatedness, leaf defences and herbivory in natural forest stands of pedunculate oak (*Quercus robur*). For this, we genotyped 703 trees from 15 stands and quantified the concentration of leaf phenolic compounds and herbivory by leaf-chewing insects at the intermediate and upper canopy layer for a subset of 235 trees. Specifically, we addressed the following questions: (i) To what extent do leaf phenolics and insect leaf herbivory vary among stands, among trees within stands and between canopy layers within trees? (ii) Do leaf phenolics and herbivory show a genetic signal when accounting for their scale-dependent variation? (iii) To what extent does variation in leaf phenolics explain patterns of leaf herbivory? By addressing these questions, we aim at combining a thorough description of cross-scale patterns typical of natural systems with insights into the biological mechanisms that underlie plant-insect herbivore relationships in non-experimental contexts.

## Material and Methods

### Study system

We performed this study in the Landes de Gascogne region (SW France) about 40 km South from Bordeaux (44°41’N, 00°51’W). The area is dominated by extensive maritime pine (*Pinus pinaster* Ait.) plantations with scattered small stands of broadleaf forest. These are usually dominated by pedunculate oak and contain other tree species like birch (*Betula pendula* L.), Pyrenean oak (*Quercus pyrenaica* Willd.), holly (*Ilex aquifolium* L.) or willows (*Salix* spp.) in minor abundance. Such stands are not subjected to intensive forest management and many are actively expanding (Gerzabek *et al.* 2017), favoured by a recent change in silvicultural management that tends to conserve oaks recruiting within adjacent pine plantations in order to increase biological pest management (Dulaurent *et al.* 2012). Pedunculate oak supports a large community of specialist and generalist herbivore insects in these stands (Giffard *et al.* 2012). Leaf chewers, skeletonizers, miners and gallers are the principal guilds responsible for background herbivory (damage imposed by a community of herbivores whose populations are at normal low densities) that amounts to values around 17.8 % (Giffard *et al.* 2012).

### Forest stands, sampling and herbivory measurements

We selected 15 forest stands of variable size and connectivity within the landscape. All stands were second-growth forests that have established since the 1950s through natural tree regeneration (Valdés-Correcher *et al.* 2019). They were strongly dominated by pedunculate oak and contained a variable but often rather sparse woody understory vegetation. The number of established oak trees ranged from 16 to 124 individuals and their surface (as derived from the minimum polygon including all trees) from 0.04 to 0.5 ha. Further information can be found in Table A1 of the Supplementary Material (see also Valdés-Correcher et al., 2019). Within each stand, we mapped and tagged every oak tree with a diameter at breast height (dbh) >3 cm and collected leaf material that was stored in silica gel until DNA isolation for the genotyping. This exhaustive sampling included a total of 703 individuals (see Table A1).

In September 2018, we randomly selected 16 individuals with a dbh >6 cm within each stand (total *n* = 235). On each tree, we haphazardly choose and cut two south-facing branches situated at 4 and 8 m above ground level, respectively, which corresponds to the intermediate (shaded) and upper (sun-exposed) tree canopies in most of our trees (see also Castagneyrol et al., 2019). Three of the 235 sampled trees did not reach 8 m, so we shifted the position of the intermediate and upper tree canopy layers 2 m downward (i.e., 2 and 6 m, respectively). Operators unaware of the study design systematically picked the 30 most apical leaves from each branch, resulting in a total of 60 leaves per tree. Samples were stored at −18°C until insect herbivory measurement (see below). For each leaf, we visually estimated the percent leaf area removed by chewing insects using the following scale: 0 = 0%, A = 1-5%, B = 6-15%, C = 16-25%, D = 26-50%, E = 51-75%, F= >75%). We used pre-established templates mimicking known levels of insect herbivory on oak leaves to increase reliability and repeatability of herbivory measurements. Herbivory levels were always estimated by the same observer (A. Bourdin) blind to leaf origin to maximise consistency of the estimates and reduce unconscious bias. We averaged values across all leaves to obtain a mean value per branch, and then used the median of each percentage class for statistical analyses (Castagneyrol *et al.* 2019).

We also collected 10 fully expanded leaves with no signs of herbivory or pathogen infection from each branch for quantification of phenolic compounds. We immediately oven-dried these leaves for 48-72 h at 45°C and grounded them to a thin powder before further chemical analyses (see below).

### Molecular analyses

Genomic DNA was isolated from the leaves using the Invisorb® DNA Plant HTS 96 kit/C and the standard protocol. All trees were genotyped using 141 single nucleotide polymorphism (SNP) markers from the sets described in Gerzabek (2017) and Guichoux (2013). The list of loci is provided in Guichoux (2013). For genotyping, SNP loci were multiplexed using an iPLEX Gold kit on a MassARRAY Typer Analyzer 4.0.26.75 (Agena Biosciences) at the Genomic and Sequencing Facility of Bordeaux (France), as described in Gerzabek et al. (2017). High-quality data with a low proportion of missing calls were obtained for all markers and individuals.

### Chemical analyses

We extracted phenolic compounds from 20 mg of dry leaf tissue with 1 mL of 70% methanol in an ultrasonic bath for 20 min, followed by centrifugation (Moreira *et al.* 2014). Samples were centrifuged at 3500 rpm and transferred to chromatographic vials. We performed the chromatographic analyses in an Ultra-High-Performance Liquid-Chromatograph (UHPLC Nexera LC-30AD; Shimadzu Corporation, Kyoto, Japan) equipped with a Nexera SIL-30AC injector and a SPD-M20A UV/VIS photodiode array detector.

For the compound separation, we used a Kinetex™ 2.6 µm C18 82-102 Å, LC Column 100 × 4.6 mm (Phenomenex, Torrance, CA, USA), protected with a C18 guard cartridge. The flow rate was established at 0.4 mL min^−1^ and the oven temperature was set at 25 ºC. The mobile phase consisted of two solvents: water-formic acid (0.05%) (A) and acetonitrile-formic acid (0.05%) (B), starting with 5% B and using a gradient to obtain 30% B at 4 min, 60% B at 10 min, 80% B at 13 min and 100 % B at 15 min. The injection volume was 15 µL. We recorded chromatograms at 330 nm and processed data with the LabSolutions software (Shimadzu Corporation, Kyoto, Japan). For phenolic compound identification, we used an ultra-performance liquid chromatography coupled with electrospray ionization quadrupole (Thermo Dionex Ultimate 3000 LC; Thermo Fisher Scientific, Waltham, MA, USA) time-of-flight mass spectrometry (UPLC-Q-TOF-MS/MS; Bruker Compact™, Bruker Corporation, Billerica, MA, USA). We identified four groups of phenolic compounds: flavonoids, ellagitannins and gallic acid derivatives (“hydrolysable tannins” hereafter), proanthocyanidins (“condensed tannins” hereafter), and hydroxycinnamic acid precursors to lignins (“lignins” hereafter). We quantified flavonoids as rutin equivalents, condensed tannins as catechin equivalents, hydrolysable tannins as gallic acid equivalents, and lignins as ferulic acid equivalents (Moreira et al., 2018; Galmán et al., 2018). We achieved the quantification of these phenolic compounds by external calibration using calibration curves at 0.25, 0.5, 1, 2 and 5 μg mL^−1^. We calculated total phenolics for each branch as the sum of flavonoids, lignins, condensed tannins and hydrolysable tannins, and expressed concentrations of each phenolic group in mg g^−1^ tissue on a dry weight basis.

### Statistical analyses

Prior to the analysis of genetic relatedness, we first examined the landscape-scale genetic structure of our oak stands by calculating pairwise *F*_st_ between stands according to Weir and C. Cockerham (1984). Overall low values (mean *F*_st_ = 0.041; range = 0.006-0.111) (**Table A2**), confirmed that the 15 stands can be considered a single gene pool and confounding effects due to population genetic structure are negligible. Then, we quantified the level of genetic relatedness between each pair of trees relative to the full sample (*n* = 703). For this, we computed a kinship matrix using Nason’s kinship coefficient (Loiselle *et al.* 1995) with the software SPAGeDi version 1.2 (Hardy, Olivier J.; Vekemans 2002). We extracted the values for our 16 target trees per stand from the global matrix and used this information as a quantitative estimate of their genetic relatedness in the subsequent analyses (Van Horn *et al.* 2008). Note that kinship-based estimates of relatedness, while commonly used in population genetics (Pemberton 2008), are not directly comparable to those obtained through direct pedigree analyses.

We modelled patterns of insect leaf herbivory and leaf phenolics at the whole-tree and at the branch level by means of linear mixed-effect models (LMM). At the tree level, we built two independent LMM with stand ID and the kinship values of the target trees as random factors in order to estimate the variance and the percentage of the overall variance explained by the local environment (stand ID) and by the genetic relatedness among trees (the kinship matrix). The first model was an intercept only model with (tree-level mean) concentration of leaf phenolics as response variable (Eq. 1). The second model included (tree-level mean) insect herbivory as response variable and leaf phenolics as an additional explanatory variable (Eq. 2).

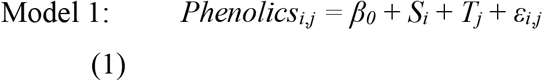

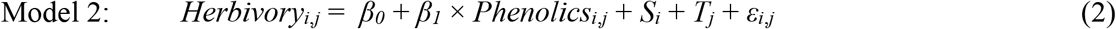

where *β*_*0*_ is the model intercept, *β*_*1*_ the fixed effects of leaf phenolics, *S*_*i*_ the random effect of stand *i*, *T*_*j*_ the random effect of tree genetic similarity *j* (entered as the kinship matrix) and *ε*_*i,j*_ the error, with *S*_*i*_ ∈ *N*(0, *σ*_*S*_^2^), *T*_*j*_ ∈ *N*(0, *σ*_*T*_^2^) and *ε*_*i,j*_ ∈ *N*(0, *σ*_*E*_^2^). For each model, we computed the variance of the fixed effects (if any, *σ*_*F*_^2^) and calculated the percentage of variance explained by each random factor (*e.g.*, 100 × *σ*_*T*_^2^ / (*σ*_*S*_^2^ + *σ*_*T*_^2^ + *σ*_*E*_^2^) for the random effect of tree genetic similarity in model 2).

The second group of models, performed at canopy level, adopted the same approach with two independent LMMs modelling the response of total leaf phenolics and herbivory, respectively (Eqs. 3 and 4):

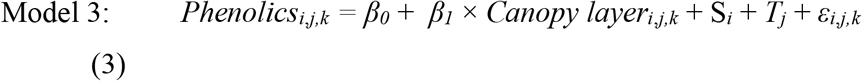

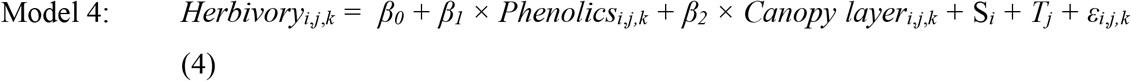

where *k* indicates the branch, *β*_0_ the intercept, *β*_1_ and *β*_2_ the coefficient parameters of the fixed effects and *S*_*i*_, *T*_*j*_ and *ε*_*i,j,k*_ as above. Again, we computed the variance of the fixed effects (if any, *σ*_*F*_^2^) and calculated the percentage of variance explained by each random factor.

All analyses were done in R version 3.5.2 (R Core Team 2018). LMMs including a kinship matrix as random factor were fit with the function *lmekin* in package *coxme* (Terry M. Therneau 2018).

## Results

### Leaf phenolics and insect herbivory at tree level

Leaf phenolic concentration was on average (± se) 14.69 ± 0.39 mg·g^−1^ (Figure 1). The random factors collectively explained 26.9 % of the overall variation, with stand ID accounting for 19.7 % and tree genetic relatedness for 7.1 %. Insect leaf herbivory was on average 12.27 ± 0.29 % and decreased significantly with increasing leaf phenolic concentration (model coefficient parameter estimate: −0.12 ± 0.05, *z* = −2.48, *P* = 0.013). The effect size was however small (2.0 %). Stand ID explained 38.1 % of the overall variation in herbivory and genetic similarity among trees only accounted for another 2.9 %.

**Figure 1.**
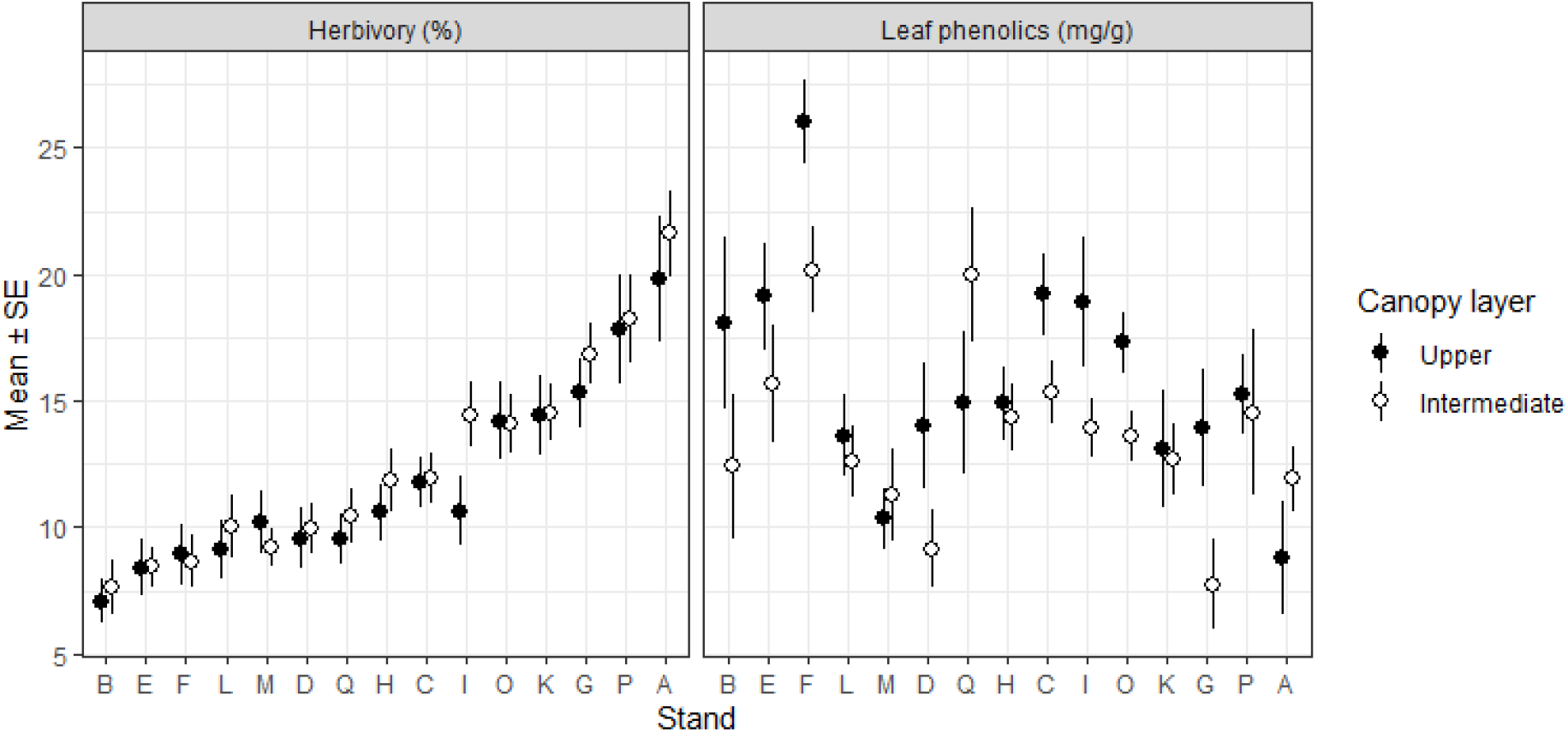
Percentage of insect herbivory and concentration of total leaf phenolics (mg/g) in the intermediate (white dots) and upper (black dots) tree canopy. Dots and error bars represent means ± SE aggregated at the level of oak stands (A-Q). Note that stands were ordered according to mean insect herbivory and the same order was used to display stand-level variability in leaf phenolics.

### Leaf phenolics and insect herbivory at canopy level

Leaf phenolic concentration was significantly lower in the intermediate than in the upper canopy layer (mean ± SE: 13.63 ± 0.50 *vs* 15.79 ± 0.59 mg·g^−1^) (Figure 1, Table 1). Stand ID accounted for 13.7 % and tree genetic relatedness for 13.9 % of the overall variation (Table 1). Insect leaf herbivory did not differ significantly between tree canopies (12.53 ± 0.39 % *vs* 11.99 ± 0.43 %) and was independent of leaf phenolic concentration (Table 1). Stand ID and tree genetic relatedness accounted for 32.0 and 34.7 % of the overall variability in insect herbivory, respectively (Table 1).

**Table 1.**
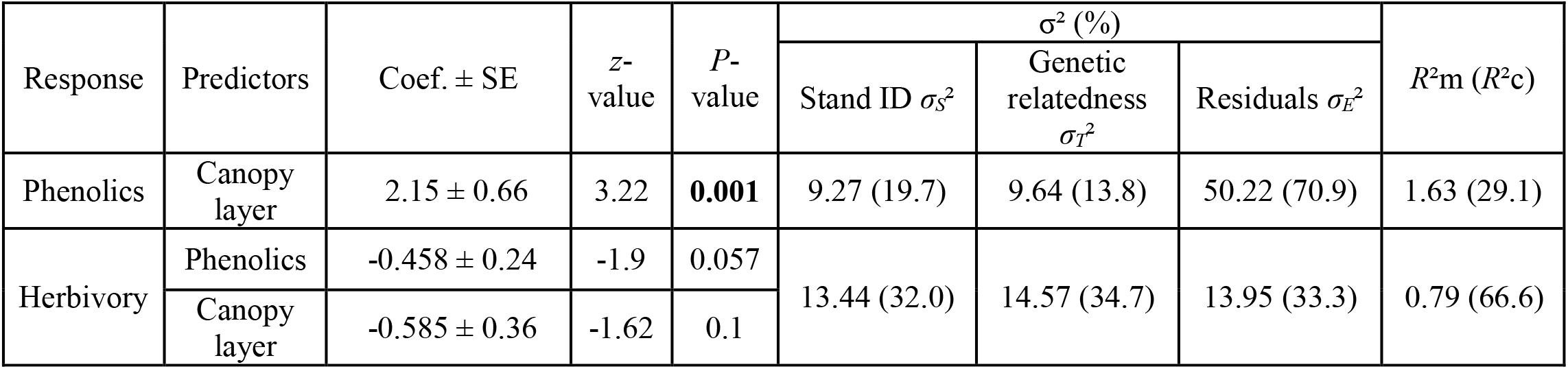
Summary of LMM testing the effect of canopy layer (upper vs. intermediate) on either leaf phenolics or insect herbivory. For insect herbivory, the effect of leaf phenolics was also included in the model. Significant variables are indicated in bold. *σ²* and % correspond to the variance and the percentage of variance explained by the random factors: stand ID, genetic similarity introduced as a kinship matrix, and the residuals. *R*²m and *R*²c correspond to the variance explained by fixed and fixed plus random factors, respectively.

| Response | Predictors | Coef. ± SE | z-value | P-value | $\sigma^2$ (%) | | | $R^2m$ ($R^2c$) |
| --- | --- | --- | --- | --- | --- | --- | --- | --- |
| | | | | | Stand ID $\sigma_s^2$ | Genetic relatedness $\sigma_I^2$ | Residuals $\sigma_e^2$ | |
| Phenolics | Canopy layer | $2.15 \pm 0.66$ | 3.22 | **0.001** | 9.27 (19.7) | 9.64 (13.8) | 50.22 (70.9) | 1.63 (29.1) |
| Herbivory | Phenolics | $-0.458 \pm 0.24$ | -1.9 | 0.057 | 13.44 (32.0) | 14.57 (34.7) | 13.95 (33.3) | 0.79 (66.6) |
| | Canopy layer | $-0.585 \pm 0.36$ | -1.62 | 0.1 | | | | |

In the intermediate canopy layer, stand ID and genetic relatedness accounted for 13.5 % and 0.03 % of the overall variation in leaf phenolics, respectively. Leaf phenolics had no significant effect on herbivory (Figure 2). Stand ID explained 40.5 % of the overall variation in herbivory, while tree genetic relatedness accounted for less than 0.02 %.

**Figure 2.**
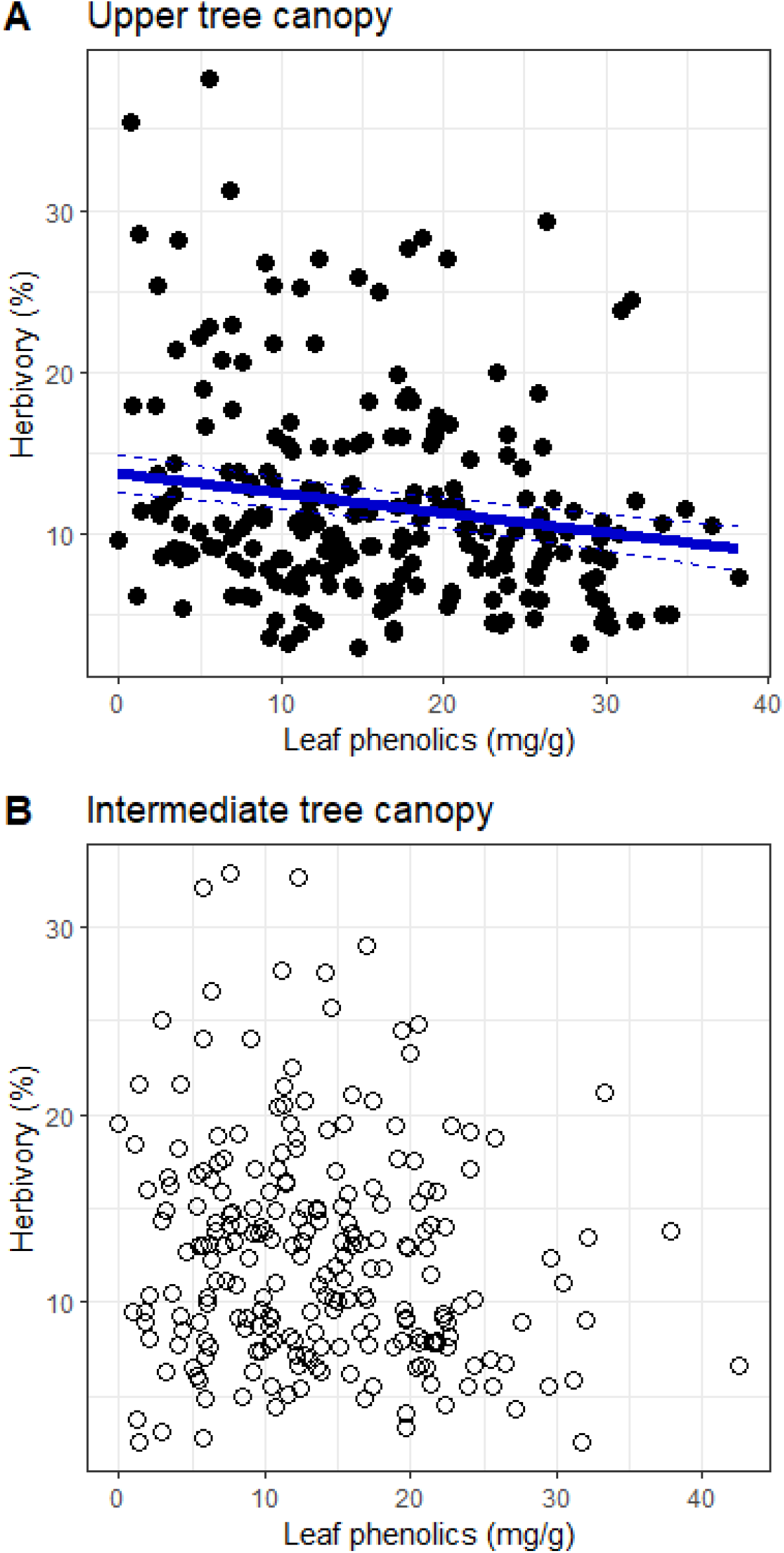
Effects of total leaf phenolics on insect herbivory in the upper (A) and intermediate (B) canopy layer. Dots represent individual trees. The thick solid line and the thin dashed lines in graph A represent model predictions and corresponding standard errors, respectively.

In the upper canopy layer, stand ID and genetic relatedness explained 17.4 % and 24.8 % of the overall variation in leaf phenolics, respectively. There was a significant, albeit weak, negative effect of leaf phenolic concentration on herbivory (coefficient parameter estimate ± SE: −0.12 ± 0.05, *z* = −2.63, *P* = 0.009) (Figure 2). Leaf phenolics accounted 2.8 % of the overall variation in herbivory while stand ID and tree genetic relatedness accounted for 25.3 and 14.5 %, respectively.

## Discussion

Tree genetic relatedness explained a noteworthy part of the overall variation in leaf phenolics and associated insect leaf herbivory. However, this genetic effect was only evident in the upper tree canopy where concentrations of leaf phenolics were consistently higher. To our knowledge, our work represents one of the first evidence of genotype-phenotype-herbivory links in natural tree populations and argues for increased consideration of canopy effects to improve our understanding of ecological and evolutionary factors driving plant-herbivore interactions on long-lived plants.

Oak trees lost between 7 and 22% of their leaf area to insect herbivores, a range of defoliation similar to previous estimates (Giffard *et al.* 2012; Castagneyrol *et al.* 2019; Valdés-Correcher *et al.* 2019). Our analysis at the whole-tree level attributed most of the overall variation in leaf herbivory to differences among forest stands whereas the contributions of tree genetic relatedness and leaf phenolics were very weak. This result might suggest that insect leaf herbivory in our system would be basically driven by the nature of the forest stand, which encapsulates diverse environmental drivers acting at the local (e.g. stand size, tree density and species composition, vegetation structure; Fuller et al., 2018; Maguire et al., 2016; van Schrojenstein Lantman et al., 2018) to landscape (e.g. stand connectivity, nature of matrix habitats; Morante-Filho et al., 2016) scale. Valdés-Correcher et al. (2019) actually reported for the same study stands that their size and connectivity affected patterns of herbivory by different insect guilds. Limiting our analyses to the whole-tree level would hence have led to the conclusion that insect herbivory is primarily determined by extrinsic drivers and shaped by the ecological neighbourhood of the focal tree.

While tree genetic relatedness had little effect on herbivory (2.9%), it was somewhat more influential in the case of leaf phenolics (7.1%) (Figure 3). Together with the likewise weak but statistically significant negative association between leaf phenolics and herbivory, one might argue that our results mirror - albeit extremely faintly - experimental studies that have consistently identified plant chemistry as the phenotypic link between the host plant genotype and the structure of associated arthropod communities (Bangert *et al.* 2006; Barbour *et al.* 2009, 2015) or patterns of herbivory (Bailey *et al.* 2006; Andrew *et al.* 2007; Donaldson and Lindroth 2007). But consistent empirical support for this linkage from natural populations remains very scarce. In one of the few available studies, Kagiya et al. (2018) found that genetic relatedness of alder (*Alnus hirsuta*) trees largely determined associated arthropod communities, yet the effect was stronger for herbivore enemies (i.e., predators) than for herbivores. Maldonado-López et al. (2015) observed that tree genetic relatedness of *Q. castanea* trees was significantly associated with chemical defences but not with insect herbivory. In turn, Tack et al. (2012) and Gossner et al. (2015) failed to detect relationships between tree genetic relatedness and herbivory in *Q. robur* populations and concluded that genetic effects tend to be overwhelmed by environmental and spatial factors.

**Figure 3.**
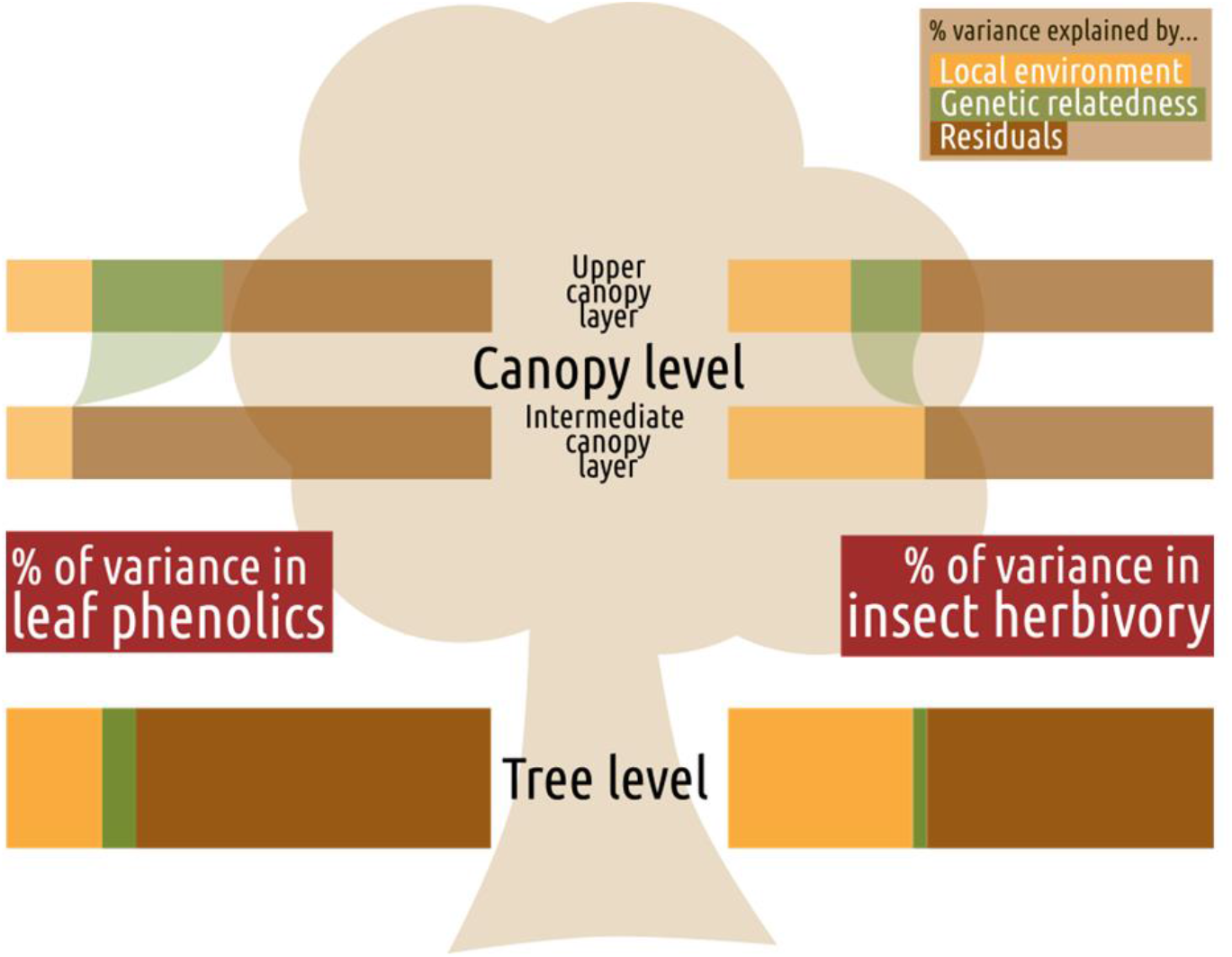
Summary of the variance partitioning among random effects and model residuals for leaf phenolics (left) and insect herbivory (right). Box length is proportional to the percentage of variance explained by each component.

In line with the predominant trend reported in the literature (e.g., Lämke and Unsicker, 2018; Poorter et al., 2006; Yamasaki and Kikuzawa, 2003; but see Roslin et al., 2006), we observed that upper canopy leaves systematically contained higher concentrations of leaf phenolics than those from the intermediate canopy. Extensive within-individual variation in leaf morphological and chemical traits is an inherent feature of plants (Niklas *et al.* 2009; Herrera 2017). For leaf phenolics, the phenomenon has been primarily explained as an ecophysiological, enzymatic and transcriptomic consequence of the higher irradiance that upper-canopy leaves receive, given that diverse phenolic compounds are involved in the protection from UV-B damage (reviewed in Jenkins and Brown, 2018); see also Lämke and Unsicker, 2018). This vertical gradient in leaf phenolics could have important consequences for plant-insect herbivore interactions. Herbivores tend to forage preferentially on upper-canopy leaves owing to their higher nutritive value (Fortin and Mauffette 2002; Oishi *et al.* 2006), yet field surveys typically report higher levels of leaf removal in lower canopy layers (e.g. Castagneyrol et al., 2019; Stiegel et al., 2017; Yamasaki and Kikuzawa, 2003), which is in line with a higher abundance and diversity of herbivores in these layers (reviewed in Ulyshen, 2011). Numerous studies have assessed within-individual variation in leaf traits and associated herbivory over the past twenty years (Lämke and Unsicker 2018), giving rise to the hypothesis that variance in nutritional quality itself could act as a defence mechanism that reduces insect herbivore performance by forcing herbivores to actively forage for suitable food (e.g. Wetzel et al., 2016; Wetzel and Meek, 2019). Yet few if any studies have addressed the implications of this within-individual variability for genotype-phenotype-herbivory relationships.

The effect of tree genetic relatedness on insect herbivory and leaf phenolics was contingent on the canopy layer. Effects were considerable in the upper canopy but negligible in the lower canopy. Sun leaves are far more productive in terms of carbon fixation than shade leaves (Poorter *et al.* 2006) and their defence against herbivores is therefore disproportionately important for overall tree performance. Our finding that tree genotypes with high phenolic compound contents in the upper canopy systematically experience lower herbivory hence suggests that such genotypes could have a non-negligible fitness advantage. On the other hand, the extent of intra-individual variability in phenolic compounds can also be heritable (Herrera 2017) and might act as an indirect defensive trait (Wetzel *et al.* 2016). If this were the case in our study system, we would expect that trees with large differences in defence allocation between upper and lower canopy leaves would tend to experience reduced herbivory. Our data did however not confirm such a trend (results not shown), suggesting that the strength of within-individual variation in leaf defences either lacks a genetic basis or has no effect on (tree-level) herbivore activity. Finally, the genetic signal in leaf herbivory that we detected suggests that leaf defences may differentially drive herbivory community heritability across different parts of the canopy. The phenomenon has been thoroughly documented at the whole-plant level in common garden experiments (e.g., Andrew et al., 2007; Robinson et al., 2012), whereas studies in natural populations have reported lower or non-significant levels of genetic variation and heritability. One important reason may be that most previous studies investigating the role of tree genetics on defences and associated herbivory have not explicitly addressed the role of the canopy layer (but instead pooled leaf samples from different heights; e.g. Gossner et al., 2015; Kagiya et al., 2018; Maldonado-López et al., 2015). Our study shows, however, that not taking within-individual variability in herbivory and defences properly into account can easily mask effect of genetic signals. Based on our findings, we recommend that future studies adopt hierarchical sampling designs and properly consider within-individual variability in both plant traits and insect herbivory when exploring their genetic basis in real-world contexts. Finally, we also recommend that further studies include other defence traits (e.g. physical defences such as trichomes and toughness or indirect defences such as volatile organic compounds) and strategies (e.g. induced defences or tolerance). Distinguishing between all these traits or strategies would allow to fully characterize multivariate defensive phenotypes (i.e. syndromes) and to better understand within and among-individual variation in genotype-phenotype-herbivory relationships.

## Acknowledgements

We are most indebted to Xavier Capdevielle, Yannick Mellerin, Inge van Halder, Gilles Saint-Jean, Victor Rébillard, Martine Martin-Clotte, and Patrick Leger for their technical assistance in the field, Gabriel Gerzabek, Erwan Guichoux, Marie Massot and Adline Delcamp for performing the molecular analyses, and Benjamin Brachi for his help with the data analyses. This work was funded by the project SPONFOREST (grant BIODIVERSA 2015-58). Genotyping was performed at the Genome Transcriptome Facility of Bordeaux (Grants from Investissements d’Avenir, Convention attributive d’aide EquipEx Xyloforest ANR-10-EQPX-16-01).

## Author contributions

E.V.C., B.C., and A.H. conceived the study. E.V.C. and A.B. acquired the data. A.B., X.M. and A.G. performed the chemical analysis. E.V.C, A.B. and B.C analysed the data with help from S.C.G.M. E.V.C. and A.H. drafted the first version of the manuscript. All authors edited the final version of the manuscript and approved its submission.

## References

Aikens KR, Timms LL, Buddle CM. 2013. Vertical heterogeneity in predation pressure in a temperate forest canopy. PeerJ 2013: 1–19.

Andrew RL, Wallis IR, Harwood CE, et al. 2007. Heritable Variation in the Foliar Secondary Metabolite Sideroxylonal in Eucalyptus ConfersCross-Resistance to Herbivores. 153: 891–901.

Bagchi R, Brown LM, Elphick CS, Wagner DL, Singer MS. 2018. Anthropogenic fragmentation of landscapes: mechanisms for eroding the specificity of plant–herbivore interactions. Oecologia 187: 521–533.

Bailey JK, Wooley SC, Lindroth RL, Whitham TG. 2006. Importance of species interactions to community heritability: A genetic basis to trophic-level interactions. Ecology Letters 9: 78–85.

Bangert RK, Turek RJ, Rehill B, et al. 2006. A genetic similarity rule determines arthropod community structure. Molecular Ecology 15: 1379–1391.

Barbour RC, Baker SC, O’Reilly-Wapstra JM, Harvest TM, Potts BM. 2009. A footprint of tree-genetics on the biota of the forest floor. Oikos 118: 1917–1923.

Barbour MA, Rodriguez-Cabal MA, Wu ET, et al. 2015. Multiple plant traits shape the genetic basis of herbivore community assembly. Functional Ecology 29: 995–1006.

Barker HL, Holeski LM, Lindroth RL. 2018. Genotypic variation in plant traits shapes herbivorous insect and ant communities on a foundation tree species. PLoS ONE 13: 1–21.

Büchel K, Fenning T, Gershenzon J, Hilker M, Meiners T. 2016. Elm defence against herbivores and pathogens: morphological, chemical and molecular regulation aspects. Phytochemistry Reviews 15: 961–983.

Bustos-Segura C, Poelman EH, Reichelt M, Gershenzon J, Gols R. 2017. Intraspecific chemical diversity among neighbouring plants correlates positively with plant size and herbivore load but negatively with herbivore damage. Ecology Letters 20: 87–97.

Carmona D, Lajeunesse MJ, Johnson MTJ. 2011. Plant traits that predict resistance to herbivores. Functional Ecology 25: 358–367.

De Casas RR, Vargas P, Pérez-Corona E, Manrique E, García-Verdugo C, Balaguer L. 2011. Sun and shade leaves of Olea europaea respond differently to plant size, light availability and genetic variation. Functional Ecology 25: 802–812.

Castagneyrol B, Giffard B, Valdés-Correcher E, Hampe A. 2019. Tree diversity effects on leaf insect damage on pedunculate oak: The role of landscape context and forest stratum. Forest Ecology and Management 433: 287–294.

Le Corff J, Marquis RJ. 1999. Differences between understorey and canopy in herbivore community composition and leaf quality for two oak species in Missouri. Ecological Entomology 24: 46–58.

Crutsinger GM, Collins MD, Fordyce JA, Gompert Z, Nice CC, Sanders NJ. 2006. Plant genotypic diversity predicts community structure and governs an ecosystem process. Science 313: 966–968.

Damestoy T, Brachi B, Moreira X, Jactel H, Plomion C, Castagneyrol B. 2019. Oak genotype and phenolic compounds differently affect the performance of two insect herbivores with contrasting diet breadth. Tree physiology 39: 615–627.

Donaldson JR, Lindroth RL. 2007. Genetics, environment, and their interaction determine efficacy of chemical defense in trembling aspen. Ecology 88: 729–739.

Dulaurent AM, Porté AJ, van Halder I, Vétillard F, Menassieu P, Jactel H. 2012. Hide and seek in forests: Colonization by the pine processionary moth is impeded by the presence of nonhost trees. Agricultural and Forest Entomology 14: 19–27.

Fernandez-Conradi P, Jactel H, Hampe A, Leiva MJ, Castagneyrol B. 2017. The effect of tree genetic diversity on insect herbivory varies with insect abundance. Ecosphere 8.

Fortin M, Mauffette Y. 2002. The suitability of leaves from different canopy layers for a generalist herbivore (Lepidoptera: Lasiocampidae) foraging on sugar maple. Canadian Journal of Forest Research 32: 379–389.

Fuller L, Fuentes-Montemayor E, Watts K, Macgregor NA, Bitenc K, Park KJ. 2018. Local-scale attributes determine the suitability of woodland creation sites for Diptera. Journal of Applied Ecology 55: 1173–1184.

Gerzabek G, Oddou-Muratorio S, Hampe A. 2017. Temporal change and determinants of maternal reproductive success in an expanding oak forest stand. Journal of Ecology 105: 39–48.

Giffard B, Corcket E, Barbaro L, Jactel H. 2012. Bird predation enhances tree seedling resistance to insect herbivores in contrasting forest habitats. Oecologia 168: 415–424.

Gossner MM, Brändle M, Brandl R, Bail J, Müller J, Opgenoorth L. 2015. Where is the extended phenotype in the wild? The community composition of arthropods on mature oak trees does not depend on the oak genotype. PLoS ONE 10: 1–16.

Guichoux E, Garnier-Géré P, Lagache L, Lang T, Boury C, Petit RJ. 2013. Outlier loci highlight the direction of introgression in oaks. Molecular Ecology 22: 450–462.

Hardy, Olivier J.; Vekemans X. 2002. Spagedi: a Versatile Computer Program To Analyse Spatial. Molecular Ecology Notes 2: 618–620.

Herrera CM. 2017. The ecology of subindividual variability in plants: Patterns, processes, and prospects. Web Ecology 17: 51–64.

Hoban S, Kelley JL, Lotterhos KE, et al. 2016. Finding the genomic basis of local adaptation: Pitfalls, practical solutions, and future directions. American Naturalist 188: 379–397.

Van Horn RC, Altmann J, Alberts SC. 2008. Can’t get there from here: inferring kinship from pairwise genetic relatedness. Animal Behaviour 75: 1173–1180.

Jenkins G, Brown B. 2018. UV-B perception and signal transduction. Annual Plant Reviews online: 155–182.

Kagiya S, Yasugi M, Kudoh H, Nagano AJ, Utsumi S. 2018. Does genomic variation in a foundation species predict arthropod community structure in a riparian forest? Molecular Ecology 27: 1284–1295.

Külheim C, Yeoh SH, Wallis IR, Laffan S, Moran GF, Foley WJ. 2011. The molecular basis of quantitative variation in foliar secondary metabolites in Eucalyptus globulus. New Phytologist 191: 1041–1053.

Lämke JS, Unsicker SB. 2018. Phytochemical variation in treetops: causes and consequences for tree-insect herbivore interactions. Oecologia 187: 377–388.

Loiselle BA, Sork VL, Nason J, Graham C. 1995. SPATIAL GENETIC STRUCTURE OF A TROPICAL UNDERSTORY SHRUB, PSYCHOTRIA OFFICINALIS (RUBIACEAE)1. 82: 1420–1425.

Maguire DY, Buddle CM, Bennett EM. 2016. Within and Among Patch Variability in Patterns of Insect Herbivory Across a Fragmented Forest Landscape. PLoS ONE 11: 1–15.

Maldonado-López Y, Cuevas-Reyes P, González-Rodríguez A, Pérez-López G, Acosta-Gómez C, Oyama K. 2015. Relationships among plant genetics, phytochemistry and herbivory patterns in Quercus castanea across a fragmented landscape. Ecological Research 30: 133–143.

McArt SH, Thaler JS. 2013. Plant genotypic diversity reduces the rate of consumer resource utilization. Proceedings of the Royal Society B: Biological Sciences 280.

Morante-Filho JC, Arroyo-Rodríguez V, Lohbeck M, Tscharntke T, Faria D. 2016. Tropical forest loss and its multitrophic effects on insect herbivory. Ecology 97: 3315–3325.

Moreira X, Mooney KA, Rasmann S, et al. 2014. Trade-offs between constitutive and induced defences drive geographical and climatic clines in pine chemical defences. Ecology Letters 17: 537–546.

Murakami M, Wada N. 1997. Difference in leaf quality between canopy trees and seedlings affects migration and survival of spring-feeding moth larvae. Canadian Journal of Forest Research 27: 1351–1356.

Murakami M, Yoshida K, Hara H, Toda MJ. 2005. Spatio-temporal variation in Lepidopteran larval assemblages associated with oak, Quercus crispula: The importance of leaf quality. Ecological Entomology 30: 521–531.

Niklas KJ, Cobb ED, Spatz HC. 2009. Predicting the allometry of leaf surface area and dry mass. American Journal of Botany 96: 531–536.

O’Rourke ME, Petersen MJ. 2017. Extending the ‘resource concentration hypothesis’ to the landscape-scale by considering dispersal mortality and fitness costs. Agriculture, Ecosystems and Environment 249: 1–3.

Oishi M, Yokota T, Teramoto N, Sato H. 2006. Japanese oak silkmoth feeding preference for and performance on upper-crown and lower-crown leaves. Entomological Science 9: 161–169.

Pemberton JM. 2008. Wild pedigrees: The way forward. Proceedings of the Royal Society B: Biological Sciences 275: 613–621.

Poorter H, Pepin S, Rijkers T, De Jong Y, Evans JR, Körner C. 2006. Construction costs, chemical composition and payback time of high- and low-irradiance leaves. Journal of Experimental Botany 57: 355–371.

Rellstab C, Gugerli F, Eckert AJ, Hancock AM, Holderegger R. 2015. A practical guide to environmental association analysis in landscape genomics. Molecular Ecology 24: 4348–4370.

Robinson KM, Ingvarsson PK, Jansson S, Albrectsen BR. 2012. Genetic Variation in Functional Traits Influences Arthropod Community Composition in Aspen (Populus tremula L.). PLoS ONE 7: 1–12.

Roslin T, Gripenberg S, Salminen JP, et al. 2006. Seeing the trees for the leaves - Oaks as mosaics for a host-specific moth. Oikos 113: 106–120.

Ruhnke H, Schädler M, Klotz S, Matthies D, Brandl R. 2009. Variability in leaf traits, insect herbivory and herbivore performance within and among individuals of four broad-leaved tree species. Basic and Applied Ecology 10: 726–736.

van Schrojenstein Lantman IM, Hertzog LR, Vandegehuchte ML, et al. 2018. Leaf herbivory is more impacted by forest composition than by tree diversity or edge effects. Basic and Applied Ecology 29: 79–88.

Stiegel S, Entling MH, Mantilla-contreras J. 2017. Reading the Leaves’ Palm: Leaf Traits and Herbivory along the Microclimatic Gradient of Forest Layers.: 1–17.

Tack AJM, Johnson MTJ, Roslin T. 2012. Sizing up community genetics: It’s a matter of scale. Oikos 121: 481–488.

Tack AJM, Ovaskainen O, Pulkkinen P, Roslin T. 2010. Spatial location dominates over host plant genotype in structuring an herbivore community Published by: Ecological Society of America Linked references are available on JSTOR for this article: Your use of the JSTOR archive indicates your acceptance of t. 91: 2660–2672.

Terry M. Therneau. 2018. coxme: Mixed Effects Cox Models. R package version 2.2-10.

Ulyshen MD. 2011. Arthropod vertical stratification in temperate deciduous forests: Implications for conservation-oriented management. Forest Ecology and Management 261: 1479–1489.

Valdés-Correcher E, van Halder I, Barbaro L, Castagneyrol B, Hampe A. 2019. Insect herbivory and avian insectivory in novel native oak forests: Divergent effects of stand size and connectivity. Forest Ecology and Management 445: 146–153.

Weir BS, C. Cockerham. 1984. Weir și Cockerham, 1984.pdf. Evolution 38: 1358–1370.

Wetzel WC, Meek MH. 2019. Physical defenses and herbivory vary more within plants than among plants in the tropical understory shrub piper polytrichum. Botany 97: 113–121.

Wetzel WC, Screen RM, Li I, et al. 2016. Ecosystem engineering by a gall-forming wasp indirectly suppresses diversity and density of herbivores on oak trees. Ecology 97: 427–438.

Wimp GM, Martinsen GD, Floate KD, Bangert RK, Whitham TG. 2005. Plant genetic determinants of arthropod community structure and diversity. Evolution 59: 61–69.

Yamasaki M, Kikuzawa K. 2003. Temporal and spatial variations in leaf herbivory within a canopy of Fagus crenata. Oecologia 137: 226–232.

